# Single cell RNA analysis of trunk neural crest cells in zebrafish identifies pre-migratory populations expressing markers of differentiated derivatives

**DOI:** 10.1101/2020.12.24.424338

**Authors:** Ezra Lencer, Rytis Prekeris, Kristin Bruk Artinger

## Abstract

The neural crest is a migratory population of stem-like cells that contribute to multiple traits including the bones of the skull, peripheral nervous system, and pigment. How neural crest cells differentiate into diverse cell types is a fundamental question in the study of vertebrate biology. Here, we use single cell RNA sequencing to characterize transcriptional changes associated with neural crest cell development in the zebrafish trunk during the early stages of migration. We show that neural crest cells are transcriptionally diverse, and identify pre-migratory populations already expressing genes associated with differentiated derivatives. Further, we identify a population of Rohon-Beard neurons that are shown to be sources of Fgf signaling in the zebrafish trunk. The data presented identify novel genetic markers for multiple trunk neural crest cell populations and Rohon-Beard neurons providing insight into previously uncharacterized genes critical for vertebrate development.

## Introduction

A fundamental question in developmental biology concerns how multipotent cell precursors produce diverse cell types (Waddington, 1957). This question is particularly relevant for the study of the vertebrate neural crest, a transient population of stem-like cells that contribute to multiple traits and form numerous cell types. Neural crest cells (NCCs) make the cartilages and bones of the skull, neurons of the peripheral nervous system, glial cells, and pigment cells among other derivatives (Anderson, 1989; Theveneau and Mayor, 2012). How and when NCCs acquire these diverse fates are long-standing questions in the study of vertebrate development, and have direct implications for both our understanding of vertebrate evolution as well as the origins of NCC derived hereditary diseases such as cleft lip and palate.

NCC development is classically conceived as a stepwise series of bifurcating cell fate decisions (Anderson, 1989; Klymkowsky et al., 2010; Simoes-Costa et al., 2014). NCCs are specified at the neural plate border, a region of ectoderm in the gastrula stage embryo that also produces cranial placodes, and in fishes and frogs, Rohon-Beard sensory neurons. Once specified, NCCs undergo an epithelial-to-mesenchymal transition, migrate throughout the embryo, and differentiate into multiple cell types. When and how NCCs differentiate into these derivative lineages remains unclear. Early studies in chick and frog suggested that migratory NCCs are multipotent, requiring cues from the migratory environment to initiate differentiation (Bronner-Fraser and Fraser, 1988; Bronner-Fraser and Fraser, 1989; Collazo et al., 1993). Other studies in zebrafish and quail suggested that NCCs are often lineage restricted prior to migration, and thus interactions among NCCs in the neural tube must somehow specify cells to different lineages before cells become migratory (Henion and Weston, 1997; Raible and Eisen, 1994; Raible and Eisen, 1996; Schilling and Kimmel, 1994)

Current understanding of the transcriptional changes associated with different stages of NCC development has largely been built from analysis of gene expression in whole tissues (Simoes-Costa et al., 2014). However, developing tissues are often transcriptionally heterogeneous, a fact highlighted by recent developments in single cell RNA sequencing (scRNA-seq) (Briggs et al., 2018; Farnsworth et al., 2020; Farrell et al., 2018; Morrison et al., 2020; Morrison et al., 2017; Saunders et al., 2019; Wagner et al., 2018). Furthermore, as cells within tissues are often at different stages of fate specification, scRNA-seq has the potential to expose transcriptional changes associated with differentiation that would have been missed by bulk sequencing experiments (Farrell et al., 2018). For instance, in chick, scRNA-seq studies of migrating cranial NCCs identified unique transcriptional signatures associated with a morphologically cryptic population of cells at the leading migratory edge (Morrison et al., 2020; Morrison et al., 2017). Thus, an emerging picture is that NCCs are more transcriptionally heterogeneous than once thought, and that a complete understanding of NCC biology must account for this heterogeneity.

Here, we use scRNA-seq to characterize the transcriptional landscape of trunk neural crest cells (tNCCs) during the early stages of migration in zebrafish. Importantly, we sample NCCs at a stage in zebrafish development when both migratory and pre-migratory tNCC populations are present in the embryo. In the trunk, NCCs migrate along one of two paths that are temporally and spatially separated. Early migrating tNCCs move between the neural tube and somitic mesoderm along what is called the medial route (Fig. 1a). These cells give rise to neurons of the peripheral nervous system, glia, and pigment cells. Later migrating tNCCs follow the dorso-lateral path between the skin ectoderm and somitic mesoderm to produce pigment cells (Fig. 1a). Thus our experiment aims to capture transcriptional changes associated with the onset of migration and the splitting of NCCs into a neuronal or pigment lineage. We show that tNCCs are a transcriptionally diverse population of cells, and that some pre-migratory tNCCs are already expressing genes associated with multiple differentiated pigment lineages. In addition we identify a unique population of Rohon-Beard neurons (RBs) and comment on the developmental similarity among RBs and NCCs. Ultimately these data create a foundation for future genetic work studying the roles of previously uncharacterized genes in the development NCCs and RBs.

**Figure 1.**
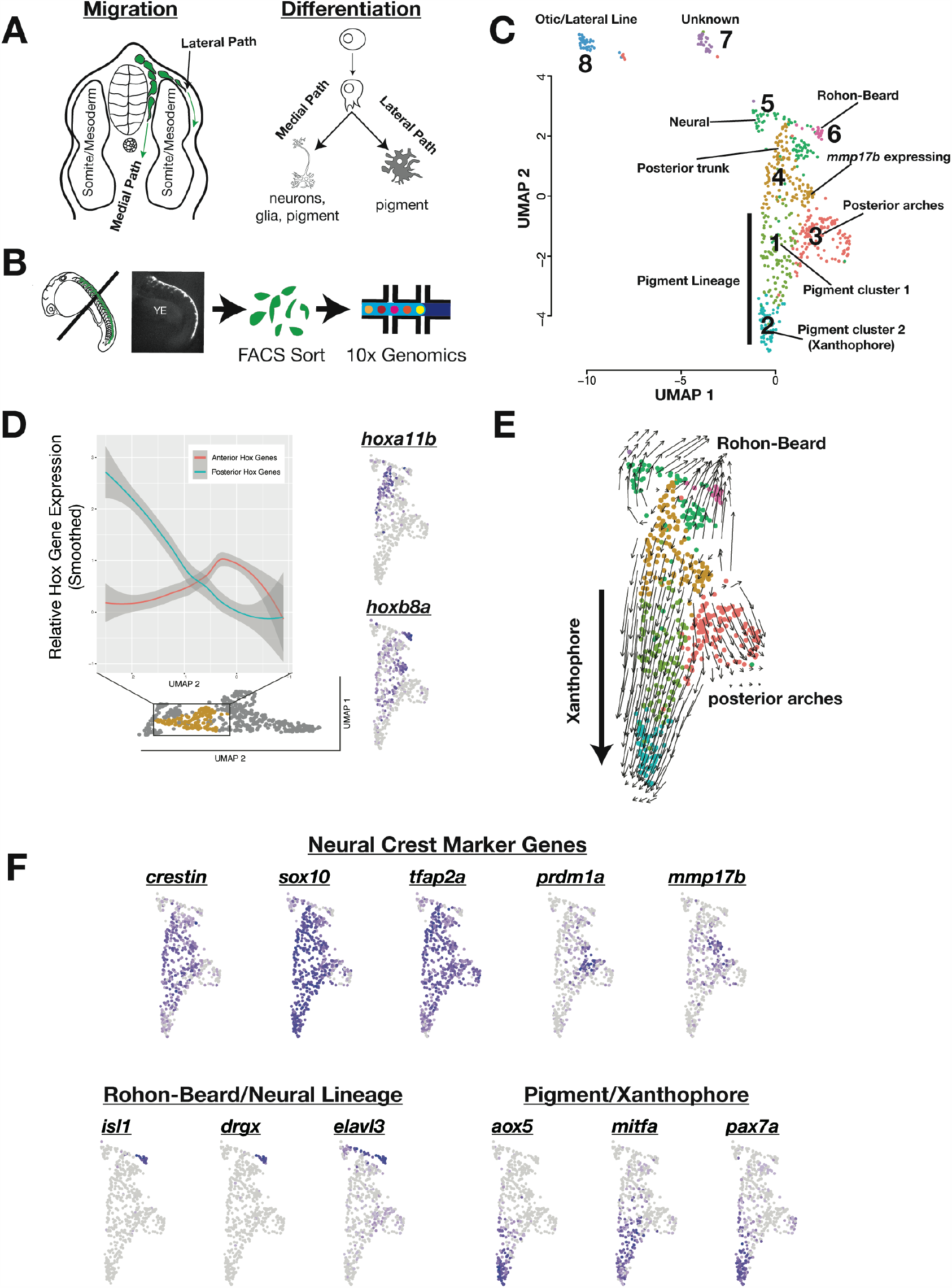
Single cell RNA-seq identifies transcriptionally heterogeneous tg(*sox10*:GFP) population. (A) Zebrafish trunk NCCs migrate along one of two routes. NCCs migrating along the medial route migrate first and produce neurons of the peripheral nervous system, glia, and pigment cells. NCCs migrating along the dorsal-lateral path produce pigment cells. (B) NCCs were sample for scRNA-seq by dissecting the trunks of tg(*sox10*:GFP) zebrafish at 24 hpf. GFP positive NCCs were FACS sorted and sequenced using 10x genomics. (C) UMAP and clustering of cells reveals multiple transcriptionally unique GFP positive cell clusters. Numbers and labels correspond to main text. (D) NCCs in cluster 4 vary in *hox* gene expression, and UMAP axis 2 broadly separates these cells along the anterior-posterior axis. Shown are smoothed anterior and posterior *hox* gene expression levels for cells in cluster 4 ordered along UMAP 2 values. Representative plots are show for anterior (*hoxb8a*) and posterior (*hoxa11b*) genes. (E) RNA velocity connects cells in developmental pseudotime. Direction and length of arrows represents predicted direction of differentiation as calculated by Velocyto (see methods). Note cells predicted to be differentiating from NCC progenitor Cluster 4 into xanthophore Cluster 2. (F) Expression of representative genes used to annotate cell clusters. Note broad expression of NCC marker genes, and localized expression of *prdm1a* marking a subset of posterior arch NC and *mmp17b* marking a subset of putative medial path migrating NCCs. RB and Xanthophore lineages are transcriptionally distinct from other cells in the dataset.

## RESULTS

### Single cell RNA-seq identifies transcriptionally unique populations of neural crest cells

We used 10x genomics to characterize the transcriptomes of 607 tg(*sox10*:GFP) positive cells from the dissected trunks of 20-24 hours post fertilization (hpf) zebrafish embryos (Fig. 1b). At this developmental stage, the tg(*sox10*:GFP) transgene labels NCCs as well as the otic organ and associated lateral line (Carney et al., 2006). Trunk NCCs (tNCCs) in the anterior segments of the body are starting to migrate along the medial pathway (Fig. 1a,b), while tNCCs that populate the posterior segments of the body, and that move along the dorso-lateral pathway, are largely pre-migratory. Thus our sample represents the early stages of tNCC migration and includes a mixed population of migratory and pre-migratory tNCCs.

Our scRNA-seq analysis identified a transcriptionally heterogeneous population of GFP positive cells. Most cells were assigned to the NCC lineage based on expression of genes known to label NCCs at this stage (*crestin, sox10*, and *tfap2a*). We also identified a cluster of cells (Cluster 8, Fig. 1c) that likely belongs to the otic organ or developing lateral line (*cldn7b, f11r*.*1, otomp, otog*), and a small cluster of cells (Cluster 7, Fig. 1c) distinguished by the expression of multiple collagen genes that we were unable to assign to a known tissue with confidence. Additionally, we find cells that likely belong to the posterior arch cranial NCCs (Cluster 3, *prdm1a, grem2b*, and *dlx2a*). Since our primary interest is in the trunk NC and associated tissues, we do not examine the otic, posterior arch, and unassigned cell clusters further.

Despite sampling zebrafish embryos during the early stages of NCC migration, we observed multiple transcriptionally unique populations of tNCCs. One cluster of tNCCs (Cluster 4) express genes that mark NCCs (e.g. *crestin, sox10, tfap2a*), and a number of Hox genes (e.g. *hoxb8a, hoxa11b, hoxb7a*) that place these cells in the posterior section of the zebrafish trunk. As NCCs in this cluster lack expression of genes identifying these cells to a differentiated lineage, we hypothesize Cluster 4 as multipotent tNCC progenitors.

A group of cells within this NCC progenitor cluster express the matrix metalloproteinase *mmp17b* (Fig. 1f), a gene known to mark tNCCs migrating along the medial pathway in the embryo (Leigh et al., 2013). To further explore this pattern, we used Seurat to sub-cluster cells that were assigned to Cluster 4 (Supplementary Fig. S2). Doing this identified that cells expressing *mmp17b* are a transcriptionally unique population, and differential expression analysis identified genes co-expressed with *mmp17b*. These include melanoma cell adhesion molecule (*mcamb)*, platelet-derived growth factor beta (*pdgfba*), small GTPase effector protein *rab11fip1a*, the kit ligand *kitlga*, and the transcription factor *prdm12b* (Supplementary Fig. S2). While *mcamb* is expressed in a salt and pepper fashion within the neural tube (Kudoh et al., 2001), both *prdm12b* and *kitlga* are expressed in regions of the embryo coincident with medial path migrating NCCs (Hultman et al., 2007; Thisse and Thisse, 2004; Zannino et al., 2014).

We further note that some cells in Cluster 4 express relatively more anterior Hox genes (e.g. *hoxc3a, hoxa4a, hoxb6a, hoxb7a, hoxb8a, hoxb9a*) and other cells express relatively more posterior Hox genes (*hoxa10b, hoxb10a, hoxd10a, hoxa11b, hoxc11b, hoxd12a*). Amazingly, the second UMAP axis broadly separates cells in Cluster 4 along this anterior-posterior Hox gene expression (Fig. 1d), suggesting that we can identify variation in our data attributable to anterior-posterior signal. Further, cells expressing *mmp17b* are inferred in this analysis as localized more anterior based on Hox gene expression. This is concordant with the known biology of tNCC migration. We sampled a developmental stage when *mmp17b* expressing tNCCs are migrating along the medial path of the more rostral body segments, while tNCCs in more caudal body segments are pre-migratory and may not yet be expressing *mmp17b*.

In addition to putative NCC progenitors, we found a large number of tNCCs already expressing genes associated with differentiated pigment cells (Fig. 1,f). This was surprising as these cells are presumably pre-migratory. Cells in Pigment Cluster 1 (Fig. 1) express melanophore/pan-pigment genes including *mitfa, gpr143, trpm1b*, among others. We hypothesize these cells as either pre-melanoblast and/or pan-pigment-cell progenitors. Cells assigned to Pigment Cluster 2 (Fig. 1c,f) express genes enriched in the xanthophore lineage (red/orange pigment) including *aox5, pax7a/b*, and *gch*. We hypothesize these cells as pre-xanthoblasts.

These data suggest that a subset of tNCCs begin expressing sets of genes associated with multiple pigment cell lineages prior to migration raising the possibility that our data reflect cells at different stages of differentiation. To explore this possibility further, we inferred RNA velocity (Velocyto) as a measure of developmental pseudotime (Bergen et al., 2020). This method uses estimates of RNA splicing to infer how cells may be transitioning across different transcriptional states. RNA velocity showed a dominant signal of tNCCs differentiating from NCC progenitor Cluster 4 into xanthophore Pigment Cluster 2 (Fig. 1e) consistent with a hypothesis that some NCCs initiate a pigment differentiation program prior to migration.

### Sox10 positive cells within the neural tube and Rohon-Beard sensory neurons are distinct from migratory neural crest

While most cells in our dataset were assigned to the NCC lineage, two cell clusters in our data (Clusters 5 and 6) are distinguished by expression of genes enriched in the neural tube and differentiated neurons (e.g. *elavl3, elavl4*, among others.). Further investigation of genes expressed by these cells suggested that these are unlikely to be NCC derived neurons. Some of these cells (Cluster 5) are presumably neural tube tissue. However, cells in Cluster 6 were transcriptionally distinct and found to express genes known to mark Rohon-Beard motor neurons including *isl1, isl2a/b*, and *drgx* (Fig. 1c,f). Rohon-Beard neurons (RBs) are a non-migratory population of cells in the dorsal neural tube that innervate the skin of embryonic and larval zebrafish. Importantly, like NCCs, RBs are also specified at the neural plate border and genes critical for the proper specification of NCCs are also critical for specification of RBs (Artinger et al., 1999; Hernandez-Lagunas et al., 2005; Hernandez-Lagunas et al., 2011). Our data recovering a putative RB population in an experiment targeting tNCCs would seem to further support some common developmental origin between these two cell types.

### Expression analysis confirms novel genetic markers in xanthophore and RB lineages

Single cell transcriptomes identify previously uncharacterized genes that are uniquely expressed in putative pre-migratory xanthoblast and RB cell populations. However, scRNA-seq cannot identify spatial expression. We thus sought to confirm expression of novel genes in the xanthophore Pigment Cluster 2 and the RBs Cluster 6 using fluorescent quantitative hybridization chain reaction (qHCR) *in-situ* hybridization (Choi et al., 2018).

#### slc2a15b and gjb8 are co-expressed in putative pre-xanthoblast tNCCs

In addition to known markers of the xanthophore lineage, our scRNA-seq data identified *slc2a15b* and *gjb8* as among the most highly expressed and restricted to the xanthophore lineage cluster (Fig.2a). The solute carrier *slc2a15b* is known to affect leucophore (white pigment cells) development in medaka (Fukamachi et al., 2006; Kimura et al., 2014; Lynn Lamoreux et al., 2005), and is expressed in the xantholeuocophore cells of adult zebrafish (Lewis et al., 2019). Leucophores and xanthophores are thought to be developmentally related (Kimura et al., 2014; Lewis et al., 2019) suggesting *slc2a15b* is a good candidate for future study. The connexin protein, *gjb8* (previously *cx30*.*3*) is known to be expressed in the skin, otic, and neural tube of zebrafish (Chang-Chien et al., 2014; Tao et al., 2010). To our knowledge no functional role for *gjb8* in NCC development has been proposed, but the importance of other gap junction proteins in pigment pattern development in zebrafish (Irion et al., 2014) makes this gene a good candidate for future study.

To confirm expression of these novel xanthophore genes to the NC we used *aox5* expression to label putative pre-xanthoblasts, and the tg(*sox10*:TagRFP) zebrafish line to mark NCCs. At 24 hpf we found that both *slc2a15b* and *gjb8* were co-expressed with *aox5* in a subset of tNCCs (Fig. 2b-f; Pearson’s r = 0.91 and 0.95 respectively). We also observed *gjb8* expression in the skin as previously reported (Tao et al., 2010), and both *gjb8* and *slc2a15b* expression in medial tissue ventral to the notochord. Within the NC, many of these *aox5/slc2a15b/gjb8* expressing cells were in the dorsal neural tube. Some were observed beginning to migrate along the dorsal lateral path and a smaller number along the medial path. Thus *aox5, slc2a15b* and *gjb8* are expressed in a subset of pre-migratory and early-migrating tNCCs. We did not observe NCCs expressing *aox5/slc2a15b/gjb8* at 19-20 hpf (20 somites) suggesting that 20-24 hpf time frame is the period during which tNCCs begin expressing markers of the xanthophore lineage.

**Figure 2.**
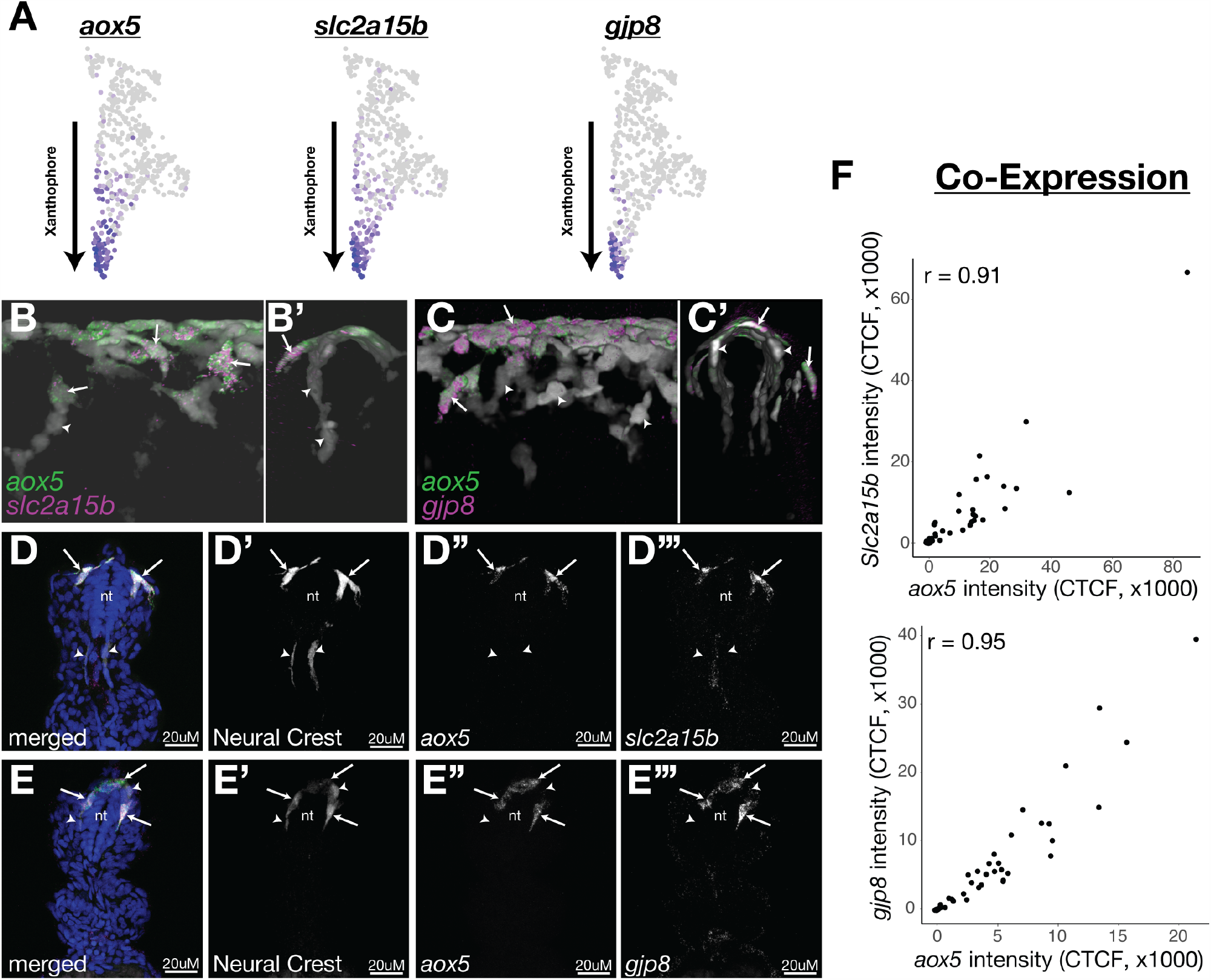
A subset of pre-migratory NCCs express known and novel marker of the xanthophore lineage. (A) Known xanthophore marker *aox5*, and two novel markers *slc2a15b and gjp8*, are predicted to be expressed in pre-migratory xanthoblast cells. (B-C) Whole mount confocal images show novel xanthophore marker genes *slc2a15b* (B) and *gjp8* (C) co-expressed in a subset of NCCs with *aox5*. Note only some NCCs co-express these genes and that most are in the dorsal neural tube. (D-E) Representative sections show co-expression of *slc2a15b* (D) and *gjp8* (E) with *aox5* in NCCs. (F) Quantification of RNA expression (intensity) in NCCs from sections shows novel markers are strongly co-expressed with *aox5*. Sections were taken every 15uM over the yolk extension. Points in panel F represent single cells taken from minimum of 3 embryos. Shown is Corrected Total Cell Fluorescence (CTCF) values (see methods). Scale bars are 20uM.

#### Rohon-Beard neurons express fgf13a/b and cxcr4b

We noticed that a number of cell signaling genes, including *fgf13a/b* and *cxcr4b*, were uniquely expressed in the putative RB cluster within our scRNA-seq data (Fig. 3a). We furthermore found these genes co-expressed in a predicted RB cell cluster from an independent scRNA-seq dataset (Wagner et al., 2018). Fgf and chemokine signaling are critical for proper morphogenesis and differentiation of numerous traits, but the expression of these developmentally important genes in RBs has not been reported before to our knowledge.

**Figure 3.**
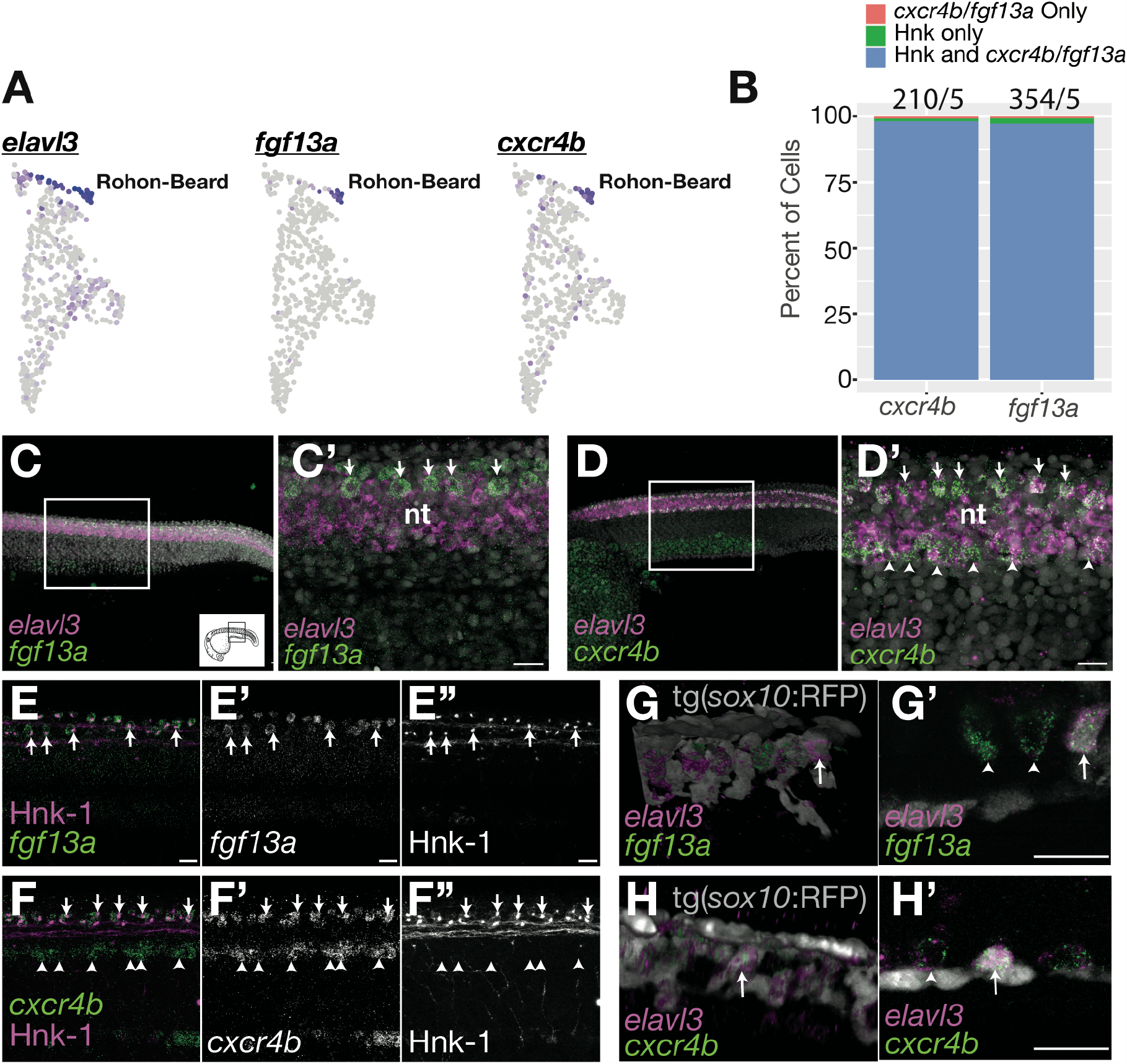
Rohon-Beard cells express *fgf13a* and *cxcr4b*. (A) Cluster of putative RBs are predicted to express neuron marker gene *elavl3* and novel RB marker genes *fgf13b* and *cxcr4b*. (B) Percent of HNK-1 positive RBs also expressing *cxcr4b* and *fgf13a* indicate that these genes are co-expressed in RB cells. The number of HNK-1, *cxcr4b*, and *fgf13a* positive cells in the dorsal neural tube were counted. Note almost all *fgf13a/cxcr4b* expressing cells co-localize to HNK-1 positive RB cells. Sample size (#cells/#embryos) reported above bars. (C-D) Representative confocal images of whole mount in situ hybridization for *elavl3, fgf13a*, and *cxcr4b* shows expression of *fgf13a* and *cxcr4b* as two rows of cells in the dorsal neural tube (arrows). *Elavl3* marks the neural tube and grey is DAPI. Note ventral neural tube expression of *cxcr4b* (arrow heads). (E) Dual in situ hybridization for *fgf13a* and immunolabeling for HNK-1 to mark RB cells shows that *fgf13a* expression overlaps with HNK-1 labeling. (F) Dual in situ hybridization for *cxcr4b* and immunolabeling for HNK-1 shows *cxcr4b expression* overlaps with HNK-1 labeling. (G-H) While *fgf13a* and *cxcr4b* expression was never observed in NCCs, a subset of putative RB cells express the tg(*sox10*:tagRFP) transgene. Shown are whole mount 3D projections (first panel) and single slice through z-stack (second panel) showing *fgf13a/cxcr4b/elavl3* positive cells co-labeled with the *sox10* transgene. Note *sox10* positive cells are positioned topologically similar to *sox10* negative RB cells. Scale bars are 20uM.

We found that f*gf13a and cxcr4b* were expressed in two rows of cells running along the lateral edges of the dorsal neural tube (*elavl3* expression) coincident with the known location of RB neurons (Fig. 3c,d). We additionally observed strong *cxcr4b* expression in the ventral neural tube (Fig. 3d), and diffuse expression of *fgf13a* in the body. Importantly, we never observed appreciable levels of *fgf13a*/*cxcr4b* expression in migrating NCCs. To confirm the identity of these *fgf13a/cxcr4b* expressing cells, we labeled RBs with HNK-1 immunofluorescence (Olesnicky et al., 2010). These data showed that dorsal trunk expression of both *fgf13a* and *cxcr4b* co-localized to HNK-1 labeled cells (Fig. 3b,e,f), indicating that these *fgf13a/cxcr4b* expressing cells are Rohon-Beard neurons. Thus RBs are apparently sources of and responsive to morphogens important for development. The function of Fgf13 ligands or the Cxcr4b receptor in RBs is unknown.

These data raise a further intriguing question: why are RB neurons present in a scRNA-seq dataset that used FACS sorting to enrich for NCCs? While we cannot rule out that RB neurons are represented in our dataset due to the proximity of RBs and NCCs to each other in the neural tube, we do note that we occasionally observe RBs labeled with the tg(*sox10*:TagRFP) transgene (Fig. 3g,h). In this zebrafish line, these RFP positive RBs were always in the same topological position in the neural tube as RFP negative RBs, indicating to us that these cells are unlikely to be NCCs. We thus hypothesize that RBs may be represented in our dataset because a small percentage of these cells express the *sox10* transgene similar to NCCs (Fig. 3, Supplementary Fig. S3). This finding is noteworthy in the context that RBs and NCCs share a developmental origin in the neural plate border (Artinger et al., 1999; Cornell and Eisen, 2000; Fritzsch and Northcutt, 1993; Hernandez-Lagunas et al., 2005). Thus our data are consistent with a literature of developmental similarity between these two cell types.

### Expression of novel xanthophore and RB neuron markers is lost in *prdm1a* mutants

To confirm expression of *slc2a15b/gjp8* in NCC derived xanthoblasts and *fgf13a*/*cxcr4b* in RBs, we examined the expression of these genes in the NRD/*prdm1a* mutant zebrafish line. Prior studies have shown that *m*utant *prdm1a*^-/-^ fish fail to specify NCCs and RBs at 24 hpf and have a reduction of melanocytes at 48 hpf (Artinger et al., 1999; Hernandez-Lagunas et al., 2005; Olesnicky et al., 2010). We thus reasoned that if expression of these genes is primarily in NCCs and RBs then we should see loss of expression of these genes in *prdm1a*^-/-^ fish and this is exactly what we found. Dorsal neural tube expression of *slc2a15b, gjp8, fgf13a*, and *cxcr4b* were all lost in *prdm1a*^-/-^ embryos (Fig. 4). Importantly, expression of these genes in areas of the embryo other than NCCs and RBs was not affected. For instance expression of *cxcr4b* in the ventral neural tube was unaffected in *prdm1a*^-/-^ embryos. These data suggest that loss of expression was due to loss of NCC ad RB cell specification, confirming expression of these genes to the NCCs and RB neurons respectively.

**Figure 4.**
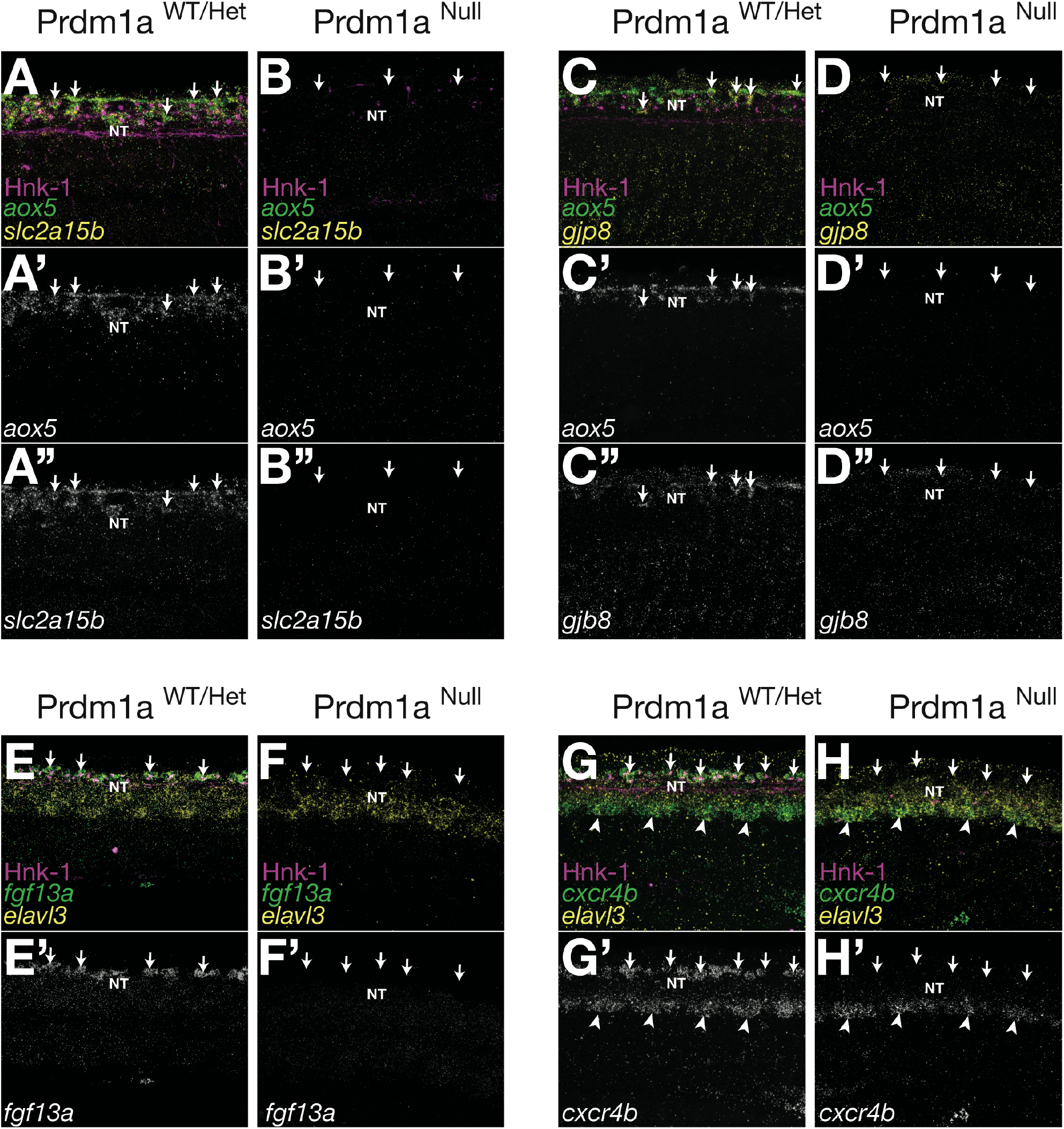
Expression of novel xanthoblast and RB neuron marker genes are lost in the Prdm1a mutant. Shown are in situ hybridizations for xanthophore (*aox5, slc2a15b, gjp8*) and RB marker genes (*fgf13a, cxcr4b, elavl3*) in wild type (A,C,E,G) and Prdm1a mutant (B,D,F,H) embryos at 24 hpf. Hnk-1 staining marks RB cells in all images. Expression of all marker genes is lost from the dorsal neural tube in Prdm1a mutant fish that lack both NCCs and RBs (arrows). Note expression of genes in regions of the embryo other than the NC and RB is still present. For instance expression of *cxcr4b* in the ventral neural tube is not altered in the Prdm1a mutant (arrow heads).

## Discussion and Conclusion

We found that tNCCs are a transcriptionally heterogeneous population during the early stages of migration, and identify a number of previously uncharacterized genes for further study. Our data indicate that a subset of NCCs begin expressing genes associated with differentiated pigment cells prior to migration and at relatively early developmental stage. In particular, cells specified towards the xanthophore lineage may be one of the earliest derivatives to begin differentiating. These data are consistent with results from cell labeling experiments in zebrafish suggesting that NCCs are lineage restricted prior to migration (Raible and Eisen, 1994; Raible and Eisen, 1996; Schilling and Kimmel, 1994). In these experiments, pre-migratory NCCs labeled in the neural tube only make a single NCC derivative. While these classic cell-labeling experiments could not identify whether pre-migratory NCCs were beginning to differentiate, our data extend these studies by showing that some pre-migratory NCCs are already expressing a pigment cell gene regulatory network. Taken together, these data support longstanding hypotheses that interactions among NCC progenitors in the neural tube are critical for specifying different NCCs towards a pigment fate (Raible and Eisen, 1996). Whether these putative pigment cells are truly fate restricted, however, is unknown and would require additional experimental manipulation.

It is worth noting that we also found a large population of putative tNCC progenitors (Cluster 4) not expressing genes associated with NCC derivatives. Thus mixed with NCCs specified towards pigment lineages are NCCs that are presumably multipotent. Some scRNA-seq studies have observed multipotent cells expressing genes associated with two or more differentiated fates (Farrell et al., 2018). These data challenge a classical bifurcating model of cell specification suggesting that instead multipotent cells begin initiating gene regulatory networks characteristic of multiple fates before becoming committed to one lineage over others. We do not observe populations of cells expressing multiple fate derivatives in our data. However, we note that our relatively small sample size for scRNA-seq datasets (607 cells) and lack of multiple sampled developmental stages may prohibit identifying these multi-fate expressing cells.

We do find a cluster of cells that we identify as putative medial path migrating tNCCs based on the expression of *mmp17b, kitlga*, and *prdm12b*. Interestingly, unlike pigment progenitors, these medial migrating tNCCs do not express genes characteristic of peripheral nervous system derivatives. Prior research in zebrafish suggests that a subset of medial migrating tNCCs act as leader cells forging a path along which follower NCCs migrate (Richardson et al., 2016). Similarly, Morrison and colleagues (2017 & 2020) used scRNA-seq to identify an unique transcriptional signature at the leading migratory edge of cranial NCCs in chick (Morrison et al., 2020; Morrison et al., 2017). Whether *mmp17b* expressing NCCs in our dataset are leader cells is uncertain. According to Leigh *et al*. 2013, *mmp17b* is not expressed in the leading medial path NCCs sugesting that these cells would not be migratory leader NCCs. Neither do we find any cluster in our data characterized by expression of orthologous genes that identify trailblazer NCCs in chick (Morrison et al., 2020; Morrison et al., 2017). Thus the identification of zebrafish trunk leader cell transcriptome remains open.

While our experimental design targeted NCCs, we also sequenced a population of RB neurons and show that in zebrafish some RBs express *sox10* and the tg(*sox10*:tagRFP) transgene. It is intriguing to consider these data in the context of other studies showing that NCCs and RBs share a similar developmental origin at the neural plate border (Artinger et al., 1999; Hernandez-Lagunas et al., 2005; Olesnicky et al., 2010). Cells at the neural plate border are thought to be an equivalence group, able to differentiate into both NCCs and RBs (Cornell and Eisen, 2002). Interactions among neural plate border cells, mediated in part by notch signaling, establish asymmetry’s that bias cells towards one fate over the other (Appel and Eisen, 2998; Cornell and Eisen, 2000, 2002). Our data suggesting RBs express both *sox10* and the tg(*sox10*:TagRFP) transgene seems to be in line with this shared developmental potential.

To what extent NCCs and RBs share a common progenitor, and whether the evolution of these two tissues is temporally coincident at the root of the vertebrate phylogeny are questions that have long captured the imagination of scholars of vertebrate biology (Artinger et al., 1999; Bone, 1960; Cornell and Eisen, 2000; Fritzsch and Northcutt, 1993). The neural crest is a novel migratory cell population unique to vertebrates with no known homology to tissues in non-vertebrate chordates (Gans and Northcutt, 1983), though see (Jeffery et al., 2008; Jeffery et al., 2004). RBs are a non-migratory neuron with uncertain homology to morphologically similar motor neurons in non-vertebrate chordates (Baker and Bronner-Fraser, 1997; Bone, 1960; Gans and Northcutt, 1983). Our data characterize the transcriptome of zebrafish RBs providing unique insight into the RB gene regulatory network. Future work identifying whether orthologous genes are expressed by motor neurons in non-vertebrate chordates could provide further insight into whether these morphologically similar neurons are homologous or homoplastic to vertebrate RB neurons (McCune and Schimenti, 2012; Wagner, 2007). Whether emerging technologies like scRNA-seq will help illuminate the evolutionary origins of new tissue types such as RBs and NCCs time will tell.

## Methods

### Zebrafish lines and husbandry

All zebrafish lines were maintained at the University of Colorado Denver | Anschutz medical school under common zebrafish husbandry conditions. Lines have been previously described and include the tg(*sox10*:eGFP)^ba2^ (Carney et al., 2006), tg(*sox10*:TagRFP)^co26TG^ (Blasky et al., 2014), and NRD^m805^/*prdm1a* mutant line (Hernandez-Lagunas et al., 2005).

### Single Cell RNA Sequencing

Fertilized Zebrafish eggs were collected in the morning, stage matched, and reared at 30C overnight until embryos were at approximately the 25 segment stage corresponding to embryos of ∼20-24 hpf at 28.5C. GFP positive embryos were dissected posterior to the otic organ using forceps in cold 1x PBS. Approximately 80 dissected trunks were pooled for dissociation and FACS sorting.

Dissected trunks were dissociated in Accumax (Innovative Cell Technologies) plus DNaseI (1uL/100uL). Cells were triturated every 15 minutes using decreasing sized pipette tips until a homogenous solution was observed. Cells were filtered through a 40uM mesh, spun, and re-suspended in Sorting buffer (1% FBS/1mM EDTA/25mM Hepes in 1x Dullbecco’s PBS (Sigma D8537)). GFP positive cells were FACS sorted on a MoFlow XDP100 cell sorter into sorting buffer and diluted to approximately 700-1100 cells/uL.

Cells were captured and prepared for sequencing using the 10X Genomics platform by the University of Colorado Denver | Anschutz Genomics Core. Libraries were sequenced on 2 lanes of an Illumina NovaSEQ6000 to an average depth of 159,520 reads per cell and a median of 666 genes per cell.

### Single Cell Analysis

Reads were mapped to the Ensembl build of the zebrafish genome (GRCz11) and UMI barcodes were assigned to unique identifiers using CellRanger (v3.2). Primary analyses were conducted using the Seurat package as implemented in R (Butler et al., 2018). Cells were filtered for quality based on mitochondrial gene expression and number of unique transcripts expressed in order to remove low quality cells as well as any possible doublets. Data was normalized using the sctransform function in Seurat while regressing out the effects of mitochondrial gene expression, ribosomal gene expression, and cell cycle (see Seurat vignettes). Cells were clustered using the first 20 PCAs and projected using the Uniform Manifold Approximation and Projection (UMAP) method as implemented in Seurat. We projected cells into 3 UMAP dimensions, but only show cells mapped onto the first two dimensions in the text for simplicity. Changing cell filtering, clustering, and UMAP projection parameters had minor affects and did not change any of the results described in the text.

Cells in Cluster 4 were further clustered in Seurat by creating a new Seurat object of only these cells and clustering under the same parameters as above.

Differential expression analysis was conducted using the Wilcox Rank Sum Test as implemented in Seurat setting the minimum fraction of cells a gene must be expressed in to 0.25 and the minimum log fold change to 0.5.

We used the package Velocyto to infer developmental pseudo-time from our data (Bergen et al., 2020). This method uses estimates of RNA splicing from scRNA-seq data to infer how cells may be transitioning across different transcriptional states; the analysis assumes that the ratio of spliced to unspliced transcripts should provide an estimate of the time since a cell has started expressing a new set of genes associated with a different transcriptional state. We used the python implementation of Velocyto to infer RNA splicing, and the R implementation of Velocyto to infer RNA velocity (e.g. pseudo-time) and plot velocities onto UMAP space.

To quantify GFP expression in RB cells, reads were aligned to the *Aequorea victoria* green fluorescence protein sequence (EBI Accession #AAA27722.1) using cell ranger (v5.0.1)

### In Situ Hybridizations and Immunofluorescence

To confirm expression of genes we used fluorescent quantitative hybridization chain reaction (qHCR) *in-situ* hybridization (Choi et al., 2018). Custom RNA probes were purchased from Molecular Instruments™. Probes were designed using either the B2 or B3 hairpin loop sequences with adaptor fluorophores of either alexafluor 488 (B3) or alexafluor 647 (B2). *In situ* hybridizations were conducted following Molecular Instruments™ protocol for whole mount zebrafish. Embryos were collected at 24hpf, fixed overnight at 4C in 4% PFA, and then stored in 100% methanol at −20C for a minimum of 24 hours. Samples were re-hydrated in PBS and permeabilized for 5 minutes in 1ug/mL proteinase K at room temperature. Hybridization was conducted using RNA probes diluted to 4 picamols/mL at 37C overnight. Amplification was performed diluting hairpins to 60 picamols/mL at room temperature overnight.

Following *in-situ* hybridization, Rohon-Beard cells were identified by HNK-1 immunofluorescence. Samples were blocked in IF blocking buffer (2% normal goat serum, 2% BSA, 1x PBS) at room temperature for a minimum of 1 hour. Primary mouse anti-HNK-1 antibody (sigma) was diluted 1:20 in blocking buffer, and samples were incubated in this primary antibody at 4C for 24-48 hours (Hernandez-Lagunas et al., 2011), and visualized using secondary goat anti-mouse IgM conjugated to alexafluor 594.

Samples were imaged on a Leica SP8 confocal microscope. For whole mount images, samples were mounted in 0.2% agarose and imaged at 40x magnification. For co-expression of xanthophore genes, samples were cryosectioned every 15uM and imaged at 63x magnification. For both whole mount and sectioned samples we imaged over the yolk extension in order to make images comparable.

Image analysis was performed using ImageJ. All images shown are 3D projections from either whole mount or sectioned material. To calculate co-expression of *aox5* with *slc2a15b* and *gjb8* we calculated Corrected Total Cell Fluorescence (CTCF) from 15uM sections. Briefly, RNA expression by a cell is proportional to relative fluorescence intensity of the HCR *in situ* signal within a cell (Choi et al., 2018). NCCs were manually segmented in 3 dimensions using the segmentation editor in ImageJ based on the expression of the tg(*sox10*:TagRFP) transgene. Average cell fluorescence intensity of *aox5*/*slc2a15b*/*gjb8*, and cell area, was calculated using the ImageJ 3D suite (Ollion et al., 2013). Average background fluorescence intensity was estimated using areas of the embryo where no signal was observed. CTCF was calculated as:

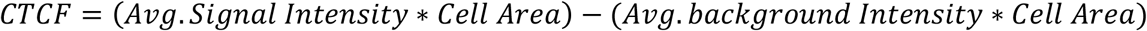

We noticed that expression of *fgf13a* and *cxcr4b* in the dorsal neural tube was more binary than quantitative. Thus we simply counted the number of HNK-1 positive cells in the dorsal neural tube that were also *fgf13a*/*cxcr4b* positive.

## Acknowledgments

We thank members of the Prekeris and Artinger laboratories for their thoughtful comments on this project and help throughout. Special thanks to Dr. Lomeli Shull for help with cell dissociation and Dr. Kenneth Jones for help discussing bioinformatics analysis. This work was funded by R01 GM122768 to RP. EL was supported by National Institutes of Health Ruth L. Kirschstein National Research Service Awards (T32CA17468 & F32HD103406). EL, RP, KA conceived of the study and wrote the manuscript. EL performed research and analyses.

## Competing Interests

The authors declare no competing interests.

## Data Availability

Data have been submitted to NCBI geo and we are awaiting an accession number. Access to geo archive will be provided prior to publication.

## Figure Legends

**Supplementary Figure S1.**
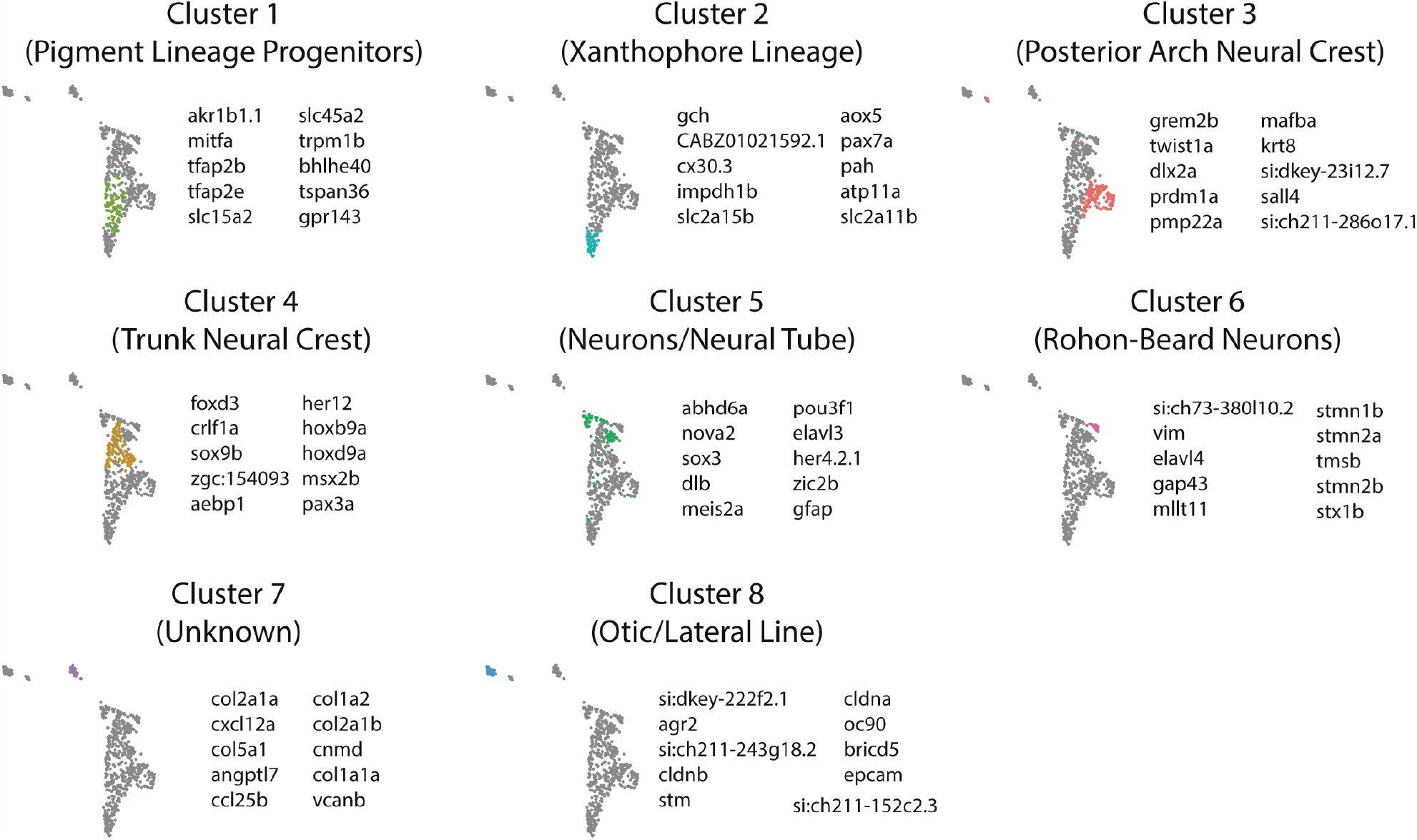
scRNA-seq clusters are identified by expression of different genes. Shown are tg(*sox10*:GFP) clusters colored onto UMAP projection. Top 10 genes that identify clusters by differential expression analysis (Wilcox rank sum) are shown for each cluster.

**Supplementary Figure S2.**
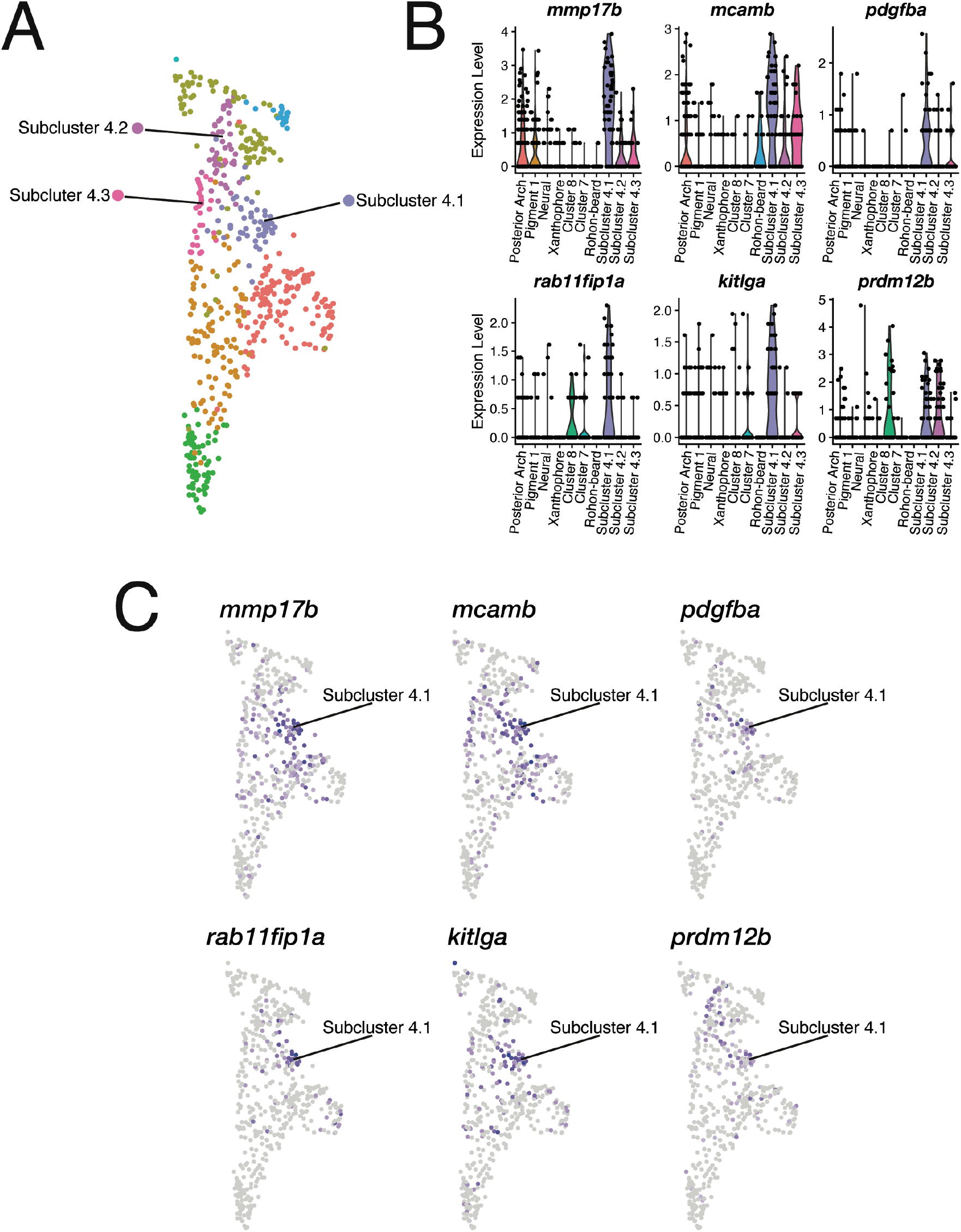
Subclustering reveals genes co-expressed with *mmp17b* in putative medial migrating tNCCs. (A) Cells in NCC progenitor cluster 4 were further separated into three subclusters based on gene expression in Seurat. Shown is UMAP with three subclusters labeled. Subcluster 4.1 are *mmp17b* expressing cells. (B) Differential expression analysis reveals genes co-expressed with *mmp17b* in subcluster 4.1 relative to all other cells. Shown are violin plots for 6 of the top genes expressed in subcluster 4.1 (see text). (C) UMAP projections show expression of 6 of the top genes co-expressed with mmp17b in subcluster 4.1.

**Supplementary Figure S3.**
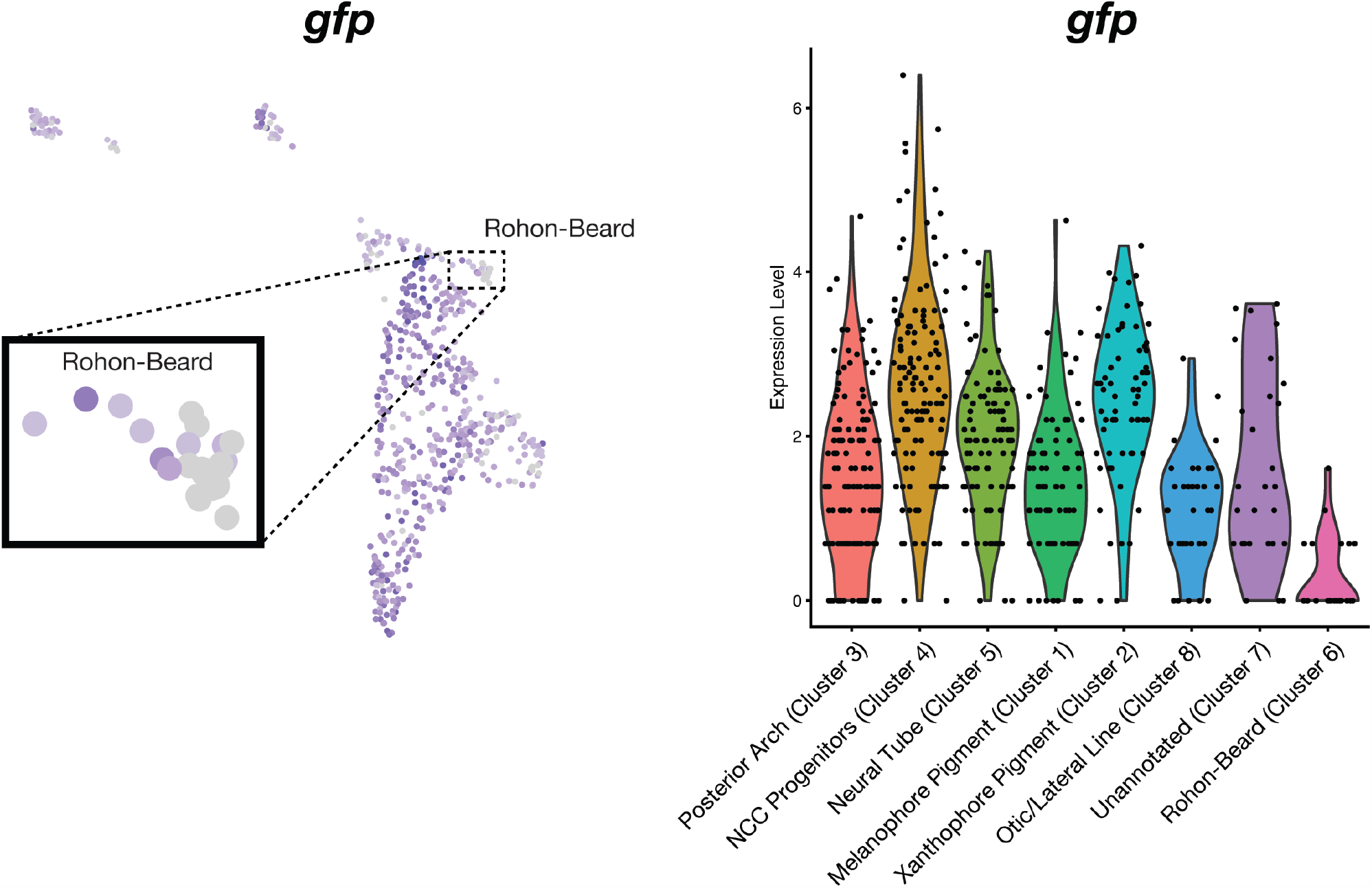
Rohon-Beard cells express low levels of the tg(*sox10*:GFP) transgene. Shown are UMAP and violin plots for the predicted expression level of eGFP transcripts in cells of each cluster. Note low level of eGFP transcripts predicted to be expressed by Rohon-Beard cells.

## Notes

### Competing Interest Statement

The authors have declared no competing interest.

## References

Anderson, D.J., 1989. The Neural Crest Cell Lineage Problem: Neuropoiesis? Neuron 3, 1–12.

Appel, B., Eisen, J.S., 2998. Regulation of neuronal specification in the zebrafish spinal cord by Delta function. Development 125, 371–380.

Artinger, K.B., Chitnis, A.B., Mercola, M., Driever, W., 1999. Zebrafish narrowminded suggests a genetic link between formation of neural crest and primary sensory neurons. Development 126, 3969–3979.

Baker, C.V.H., Bronner-Fraser, M., 1997. The origins of the neural crest. Part II: an evolutionary perspective. Mechanisms of Development 69, 13–29.

Bergen, V., Lange, M., Peidli, S., Wolf, F.A., Theis, F.J., 2020. Generalizing RNA velocity to transient cell states through dynamical modeling. Nat Biotechnol.

Blasky, A.J., Pan, L., Moens, C.B., Appel, B., 2014. Pard3 regulates contact between neural crest cells and the timing of Schwann cell differentiation but is not essential for neural crest migration or myelination. Dev Dyn 243, 1511–1523.

Bone, Q., 1960. The central nervous system in amphioxus. J. Comp. Neurol. 115, 27–51.

Briggs, J.A., Weinreb, C., Wagner, D.E., Megason, S., Peshkin, L., Kirschner, M.W., Klein, A.M., 2018. The dynamics of gene expression in vertebrate embryogenesis at single-cell resolution. Science 360.

Bronner-Fraser, M., Fraser, S.E., 1988. Cell lineage analysis reveals multipotency of some avian neural crest cells. Nature 335, 161–164.

Bronner-Fraser, M., Fraser, S.E., 1989. Developmental Potential of Avian Trunk Neural Crest Cells In Situ. Neuron 3, 755–766.

Butler, A., Hoffman, P., Smibert, P., Papalexi, E., Satija, R., 2018. Integrating single-cell transcriptomic data across different conditions, technologies, and species. Nat Biotechnol 36, 411–420.

Carney, T.J., Dutton, K.A., Greenhill, E., Delfino-Machin, M., Dufourcq, P., Blader, P., Kelsh, R.N., 2006. A direct role for Sox10 in specification of neural crest-derived sensory neurons. Development 133, 4619–4630.

Chang-Chien, J., Yen, Y.C., Chien, K.H., Li, S.Y., Hsu, T.C., Yang, J.J., 2014. The connexin 30.3 of zebrafish homologue of human connexin 26 may play similar role in the inner ear. Hear Res 313, 55–66.

Choi, H.M.T., Schwarzkopf, M., Fornace, M.E., Acharya, A., Artavanis, G., Stegmaier, J., Cunha, A., Pierce, N.A., 2018. Third-generation in situ hybridization chain reaction: multiplexed, quantitative, sensitive, versatile, robust. Development 145.

Collazo, A., Bronner-Fraser, M., Fraser, S.E., 1993. Vital dye labellin of Xenopus laevis trunk neural crest reveals multipotency and novel pathways of migration. Development 118, 363–376.

Cornell, R.A., Eisen, J.S., 2000. Delta signaling mediates segregation of neural crest and spinal sensory neurons from zebrafish lateral neural plate. Development 127, 2873–2882.

Cornell, R.A., Eisen, J.S., 2002. Delta/Notch signaling promotes formation of zebrafish neural crest by repressing Neurogenin 1 function. Development 129, 2639–2648.

Farnsworth, D.R., Saunders, L.M., Miller, A.C., 2020. A single-cell transcriptome atlas for zebrafish development. Dev Biol 459, 100–108.

Farrell, J.A., Wang, Y., Riesenfeld, S.J., Shekhar, K., Regev, A., Schier, A.F., 2018. Single-cell reconstruction of developmental trajectories during zebrafish embryogenesis. Science 360.

Fritzsch, B., Northcutt, R.G., 1993. Cranial and spinal nerve organization in amphioxus and lampreys: evidence for an ancestral craniate pattern. Acta Anat 148, 96–109.

Fukamachi, S., Wakamatsu, Y., Mitani, H., 2006. Medaka double mutants for color interfere and leucophore free: characterization of the xanthophore-somatolactin relationship using the leucophore free gene. Dev Genes Evol 216, 152–157.

Gans, C., Northcutt, R.G., 1983. Neural Crest and the Origin of Vertebrates: A New Head. Science 220, 268–274.

Henion, P.D., Weston, J.A., 1997. Timing and pattern of cell fate restrictions in the neural crest lineage. Development 124, 4351–4359.

Hernandez-Lagunas, L., Choi, I.F., Kaji, T., Simpson, P., Hershey, C., Zhou, Y., Zon, L., Mercola, M., Artinger, K.B., 2005. Zebrafish narrowminded disrupts the transcription factor prdm1 and is required for neural crest and sensory neuron specification. Dev Biol 278, 347–357.

Hernandez-Lagunas, L., Powell, D.R., Law, J., Grant, K.A., Artinger, K.B., 2011. prdm1a and olig4 act downstream of Notch signaling to regulate cell fate at the neural plate border. Dev Biol 356, 496–505.

Hultman, K.A., Bahary, N., Zon, L.I., Johnson, S.L., 2007. Gene Duplication of the zebrafish kit ligand and partitioning of melanocyte development functions to kit ligand a. PLoS Genet 3, e17.

Irion, U., Frohnhofer, H.G., Krauss, J., Colak Champollion, T., Maischein, H.M., Geiger-Rudolph, S., Weiler, C., Nusslein-Volhard, C., 2014. Gap junctions composed of connexins 41.8 and 39.4 are essential for colour pattern formation in zebrafish. Elife 3, e05125.

Jeffery, W.R., Chiba, T., Krajka, F.R., Deyts, C., Satoh, N., Joly, J.S., 2008. Trunk lateral cells are neural crest-like cells in the ascidian Ciona intestinalis: insights into the ancestry and evolution of the neural crest. Dev Biol 324, 152–160.

Jeffery, W.R., Strickler, A.G., Yamamoto, Y., 2004. Migratory neural crest-like cells form body pigmentation in a urochordate embryo. Nature 431, 696 – 699.

Kimura, T., Nagao, Y., Hashimoto, H., Yamamoto-Shiraishi, Y., Yamamoto, S., Yabe, T., Takada, S., Kinoshita, M., Kuroiwa, A., Naruse, K., 2014. Leucophores are similar to xanthophores in their specification and differentiation processes in medaka. Proc Natl Acad Sci U S A 111, 7343–7348.

Klymkowsky, M.W., Rossi, C.C., Artinger, K.B., 2010. Mechanisms driving neural crest induction and migration in the zebrafish and Xenopus laevis. Cell Adh Migr 4, 595–608.

Kudoh, T., Tsang, M., Hukriede, N.A., Chen, X., Dedekian, M., Clarke, C.J., Kiang, A., Schultz, S., Epstein, J.A., Toyama, R., Dawid, I.B., 2001. A gene expression screen in zebrafish embryogenesis. ZFIN Direct Data Submission.

Leigh, N.R., Schupp, M.O., Li, K., Padmanabhan, V., Gastonguay, A., Wang, L., Chun, C.Z., Wilkinson, G.A., Ramchandran, R., 2013. Mmp17b is essential for proper neural crest cell migration in vivo. PLoS One 8, e76484.

Lewis, V.M., Saunders, L.M., Larson, T.A., Bain, E.J., Sturiale, S.L., Gur, D., Chowdhury, S., Flynn, J.D., Allen, M.C., Deheyn, D.D., Lee, J.C., Simon, J.A., Lippincott-Schwartz, J., Raible, D.W., Parichy, D.M., 2019. Fate plasticity and reprogramming in genetically distinct populations of Danio leucophores. Proc Natl Acad Sci U S A 116, 11806–11811.

Lynn Lamoreux, M., Kelsh, R.N., Wakamatsu, Y., Ozato, K., 2005. Pigment pattern formation in the medaka embryo. Pigment Cell Res 18, 64–73.

McCune, A., Schimenti, J., 2012. Using Genetic Networks and Homology to Understand the Evolution of Phenotypic Traits. Current Genomics 13, 74–84.

Morrison, J.A., McLennan, R., Teddy, J.M., Scott, A.R., Kasemeier-Kulesa, J.C., Gogol, M.M., Kulesa, P.M., 2020. Transcriptome profiling of the branchial arches reveals cell type composition and a conserved signature of neural crest cell invasion. BioRxiv.

Morrison, J.A., McLennan, R., Wolfe, L.A., Gogol, M.M., Meier, S., McKinney, M.C., Teddy, J.M., Holmes, L., Semerad, C.L., Box, A.C., Li, H., Hall, K.E., Perera, A.G., Kulesa, P.M., 2017. Single-cell transcriptome analysis of avian neural crest migration reveals signatures of invasion and molecular transitions. Elife 6.

Olesnicky, E., Hernandez-Lagunas, L., Artinger, K.B., 2010. prdm1a Regulates sox10 and islet1 in the development of neural crest and Rohon-Beard sensory neurons. Genesis 48, 656–666.

Ollion, J., Cochennec, J., Loll, F., Escude, C., Boudier, T., 2013. TANGO: A generic tool for high-throughput 3D image analysis for studying nuclear organization. Bioinformatics 29, 1840– 1841.

Raible, D.W., Eisen, J.S., 1994. Restriction of neural crest cell fate in the trunk of the embryonic zebrafish. Development 120, 495–503.

Raible, D.W., Eisen, J.S., 1996. Regulative interactions in zebrafish neural crest. Development 122, 501–507.

Richardson, J., Gauert, A., Briones Montecinos, L., Fanlo, L., Alhashem, Z.M., Assar, R., Marti, E., Kabla, A., Hartel, S., Linker, C., 2016. Leader Cells Define Directionality of Trunk, but Not Cranial, Neural Crest Cell Migration. Cell Rep 15, 2076–2088.

Saunders, L.M., Mishra, A.K., Aman, A.J., Lewis, V.M., Toomey, M.B., Packer, J.S., Qiu, X., McFaline-Figueroa, J.L., Corbo, J.C., Trapnell, C., Parichy, D.M., 2019. Thyroid hormone regulates distinct paths to maturation in pigment cell lineages. Elife 8.

Schilling, T., Kimmel, C.B., 1994. Segment and cell type lineage restrictions during pharyngeal arch development in the zebrafish embryo. Development 120, 483–494.

Simoes-Costa, M., Tan-Cabugao, J., Antoshechkin, I., Sauka-Spengler, T., Bronner, M.E., 2014. Transcriptome analysis reveals novel players in the cranial neural crest gene regulatory network. Genome Res 24, 281–290.

Tao, L., DeRosa, A.M., White, T.W., Valdimarsson, G., 2010. Zebrafish cx30.3: identification and characterization of a gap junction gene highly expressed in the skin. Dev Dyn 239, 2627– 2636.

Theveneau, E., Mayor, R., 2012. Neural crest delamination and migration: from epithelium-to-mesenchyme transition to collective cell migration. Dev Biol 366, 34–54.

Thisse, B., Thisse, C., 2004. Fast Release Clones: A High Througput Expression Analysis. ZFIN Direct Data Submission.

Waddington, C.H., 1957. The strategy of the Genes. Allen & Unwin, London.

Wagner, D.E., Weinreb, C., Collins, Z.M., Briggs, J.A., Megason, S.G., Klein, A.M., 2018. Single-cell mapping of the gene expression landscapes and lineage in the zebrafish embryo. Science 360, 981–987.

Wagner, G.P., 2007. The developmental genetics of homology. Nature Genetics 8, 473–479.

Zannino, D.A., Downes, G.B., Sagerstrom, C.G., 2014. prdm12b specifies the p1 progenitor domain and reveals a role for V1 interneurons in swim movements. Dev Biol 390, 247–260.

